# Duet: SNP-Assisted Structural Variant Calling and Phasing Using Oxford Nanopore Sequencing

**DOI:** 10.1101/2022.07.04.498779

**Authors:** Yekai Zhou, Amy Wing-Sze Leung, Syed Shakeel Ahmed, Tak-Wah Lam, Ruibang Luo

## Abstract

**Background:** Whole genome sequencing (WGS) using the long-read Oxford Nanopore Technologies (ONT) MinION sequencer provides a cost-effective option for structural variant (SV) detection in clinical applications. Despite the advantage of using long reads, however, accurate SV calling and phasing are still challenging.

**Results:** We introduce Duet, an SV detection tool optimized for SV calling and phasing using ONT data. The tool uses novel features integrated from both SV signatures and single-nucleotide polymorphism (SNP) signatures, which can accurately distinguish SV haplotype from a false signal. Duet was benchmarked against state-of-the-art tools on multiple ONT sequencing datasets of sequencing coverage ranging from 8X to 40X. At low sequencing coverage of 8X, Duet performs better than all other tools in SV calling, SV genotyping and SV phasing. When the sequencing coverage is higher (20X to 40X), the F1-score for SV phasing is further improved in comparison to the performance of other tools, while its performance of SV genotyping and SV calling remains comparable or higher than other tools.

**Conclusion:** Duet can perform accurate SV calling, SV genotyping and SV phasing using low-coverage ONT data, making it very useful for low-coverage genomes. It has great performance when scaled to high-coverage genomes, which is adaptable to various clinical applications. Duet is open source and is available at https://github.com/yekaizhou/duet.

## Background

The importance of structural variants (SVs) in genetic disorders has been elaborated in recent studies using traditional molecular techniques [1-3]. Third-generation sequencing (TGS) generates long-reads, which improve the detection resolution down to the base-pair level and are therefore informative enough to achieve phasing [4, 5]. Resolving SV haplotypes can be useful for determining allele-specific expression, compound heterozygosity, and other diplotypic effects [6-10], thus supporting more accurate diagnosis and treatment planning. One of the more cost-effective TGS options for SV detection involves applying the Oxford Nanopore Technologies (ONT) MinION sequencer for whole genome sequencing.

However, current SV detection tools including cuteSV [11], SVIM [12], NanoVar [13], and Sniffles [14] have several constraints. First, the accuracy of SV calling can only be guaranteed when the provided sequencing coverage is high, which is not always available in actual clinical settings. The unsatisfactory performance of existing state-of-the-art tools for low-coverage sequencing data originates from their quality control methods. These tools identify false signals when the proportion of support reads with SV marks (p_SV_) is low [11-14]. This approach works well for high-coverage sequencing data where there are abundant number of support reads, and a clear gap in p_SV_ between the true SV calls and the false signals, which can be observed and used for quality control. However, in low-coverage settings, it is common that there are only one or two support reads with SV marks for both true and false signals. Low-coverage data will generate SV candidates with low and very close p_SV_, which are hard to distinguish from true SVs and false signals based on this single feature. Therefore, novel support read signatures and novel features may add to the granularity of the quality control workflow, which is promising for boosting the performance of low-coverage SV calling.

Second, these tools can only distinguish the genotype of the detected SVs (SV genotyping). The genotype can be assigned homozygous (1/1) or heterozygous (0/1) if the level of p_SV_ is high or median, but the tools lose discriminative power when the sequencing coverage is low. Moreover, this method does not distinguish the paternal (1|0) or maternal (0|1) haplotype of the heterozygous SVs (SV phasing), which is important for clinical applications. To achieve SV phasing and accurate SV genotyping in low-coverage, it is reasonable to solicit additional support read signatures, such as the haplotype tendencies of the reads potentially from single-nucleotide polymorphism (SNP) calling and phasing. The additional signatures serve as the raw materials to derive novel features, which provide the granularity for SV phasing and quality control in SV calling at low-coverage.

In this paper, we introduce Duet, a tool for SV calling, genotyping, and phasing, optimized for ONT data. Instead of relying solely on the SV signatures on the reads [11-14], Duet incorporates SNP signatures to observe paternal or maternal tendencies of each SV support read. The tool further integrates both SV and SNP signatures into several novel features. The features form as an interpretable and robust decision path, which can characterize SV haplotype from false signals, even when the number of SV support reads is moderate. Therefore, while most existing approaches for SV phasing, require both high-coverage [15] and multi-platform sequencing data [16-19], Duet can accurately call and phase SVs with only 8X whole genome sequencing (WGS) ONT data, and has great performance in scaling when the sequencing coverage goes higher, which is promising in various clinical applications.

### Implementation

The schematic diagram of Duet is depicted in Fig. 1. Taking a long-read alignment [20] and its reference as input (Fig. 1A), Duet processes the data with four major modules, comprising (1) SNP calling (Fig. 1B), (2) SNP phasing and per-read haplotype assignment (Fig. 1C), (3) SV calling (Fig. 1D), and (4) SV phasing (Fig. 1E-F) to output phased SVs. The tool is designed to tolerate false positives of low-coverage ONT WGS data while retaining a high level of accuracy, although the use of a high-coverage dataset (i.e., ≥ 20X) can enhance its performance.

**Fig. 1.**
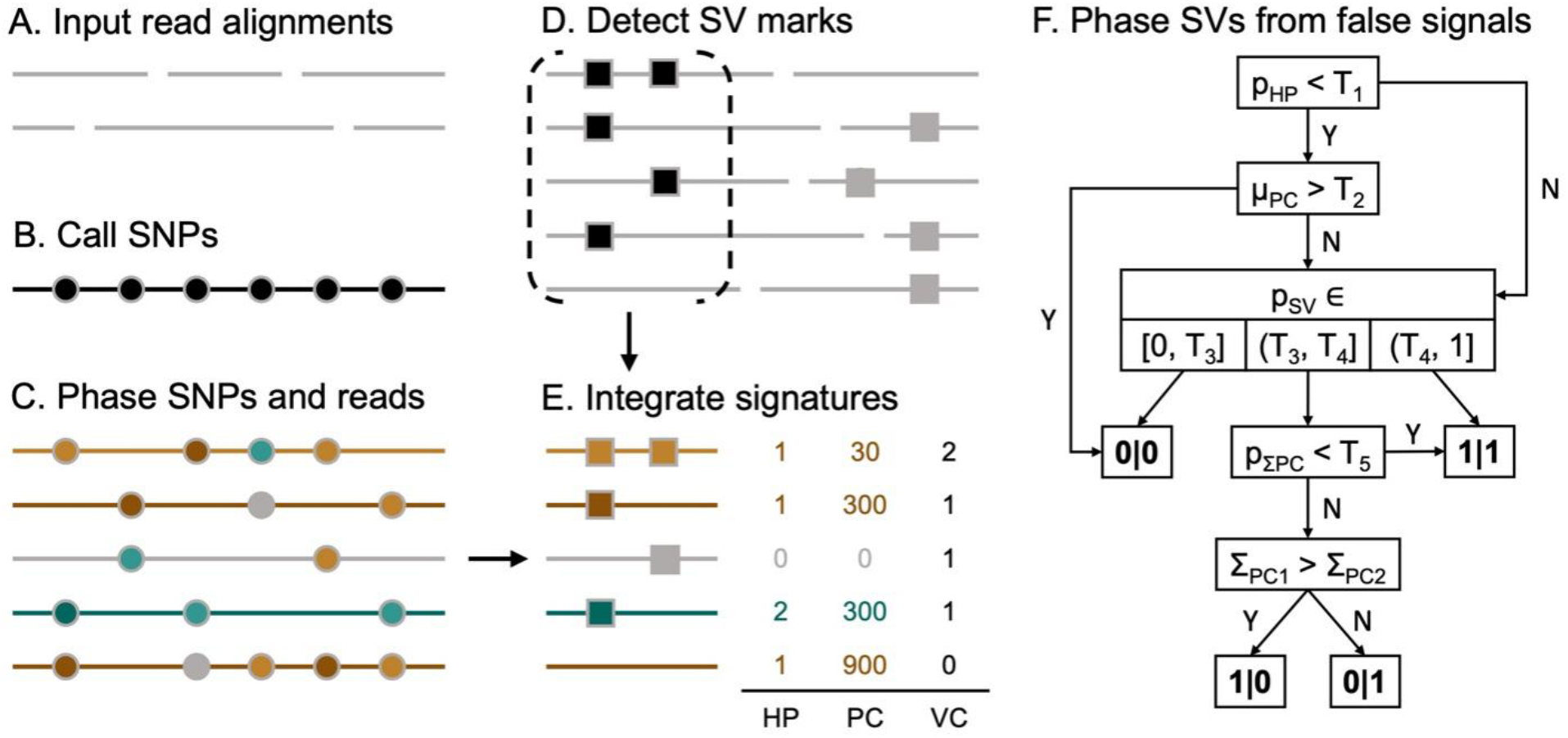
Workflow of Duet. (A) First, ONT long reads are aligned using Minimap2. (B-C) To obtain the per-read phasing information (green or brown) with its confidence level (luminance of the color), SNPs (indicated in circles) are called using Clair3 and then phased using WhatsHap. Based on the phased SNPs, the haplotypes of the reads are determined by WhatsHap. (D) The SV marks on each read are detected by SVIM. (E) Three signatures from previous steps are integrated as the signature of the support reads. (F) Duet phases SV and filters out false signals based on the features derived from the signatures in step (E). Description of the signatures and features at step (E) and step (F) is presented in Table 1. T_1_ to T_5_ are thresholds for each feature.

### SNP calling

SNP calling generates heterozygous SNPs that can be used as a proxy to infer the reads’ haplotypes. We applied the ONT model of Clair3 [21], a fast and accurate long-read small-variant caller, with “--snp_min_af 0.25 --pileup_only --call_snp_only” options after experimenting with different settings. The settings were fine-tuned in Duet, which increased the overall processing speed, in addition to providing sufficient variant-calling accuracy for phasing.

**Table 1.** Signatures and features used by Duet.

| Support read signature |  |  |
| --- | --- | --- |
| HP |  | haplotype |
| PC |  | confidence score of the haplotype prediction |
| VC |  | count of SV marks |
| Feature for SV phasing |  |  |
| $p_{SV}$ | $= \Sigma_{VC>0} / \Sigma_{VC \geq 0}$ | proportion of reads with SV marks |
| $p_{HP}$ | $= \Sigma_{VC,HP>0} / \Sigma_{VC>0}$ | proportion of phasable reads with SV marks |
| $\Sigma_{PC}$ | $= \Sigma_{VC*PC \text{ per HP}}$ | total PC for read set from each haplotype |
| $\mu_{PC}$ | $= \mu_{VC*PC \text{ per HP}}$ | average PC for read set from each haplotype |
| $p_{\Sigma PC}$ | $= \Sigma_{PC1} / \Sigma_{PC2}$ | ratio of $\Sigma_{PC}$ for both haplotype |

### SNP phasing and per-read haplotype assignment

Using the SNP-calling results, Duet applies WhatsHap [22] with “whatshap phase --distrust-genotypes” to generate the two parental haplotypes. The settings provide WhatsHap with higher tolerance for false positives, especially when working with low-coverage datasets. The haplotag “whatshap haplotag --tag-supplementary” subfunction is then called to assign haplotype to each read. In addition, each read is assigned a confidence score by WhatsHap, which is positively related to the number of accurately phased variants. Duet will use the haplotype and the prediction confidence of the reads. We employ GNU Parallel [23] to allow parallel processing of all chromosomes.

### SV calling

It is essential to apply an SV caller that is sensitive even when using low-coverage data to reduce the initial false-negative rate, as a large proportion of false positives can be filtered out after SV phasing. After performance evaluation, SVIM [12] with the setting “--minimum_depth 0 --minimum_score 0 --cluster_max_distance 0.9” provides the best output to fit the purpose of Duet, where the SV marks detected are loosely clustered for downstream analysis.

### SV phasing

Duet integrates the SV and SNP signatures obtained from the above modules to represent each support read, i.e., haplotype (HP), confidence of the haplotype prediction (phasing confidence, PC), and the number of SV marks (VC). For each SV call set with multiple support reads, several novel features are derived from the signatures, and an empirical rule-based decision path with corresponding cut-off values is derived accordingly.

The decision path contains three layers:

First, an initial filter is applied to filter out SV calls that contain too small a proportion of phasable reads (p_HP_) with a very high average phasing confidence (μ_PC_).

The remaining SV calls will go through a subsequent characterization based on the proportion of reads containing SV marks (p_SV_), to identify false positive SV calls (0/0) if with too low p_SV_, and homozygous variants (1|1) if with high p_SV_.

The further remaining SV calls with moderate proportion of reads with SV marks will be finally distinguished between being homozygous or heterozygous, where the phasing confidence is taken into meticulous consideration: Duet classifies the support reads into paternal group and maternal group based on their haplotype. Then the sum of the phasing confidence for each haplotype group is calculated (Σ_PC1_, Σ_PC2_). If these two values have no huge difference (i.e., a moderate p_ΣPC_), the SV call will be classified as homozygous (1|1). Otherwise, if there is a huge difference between the total phasing confidence, the SV call will be classified as heterozygous and assigned to the haplotype with the dominant total phasing confidence (0|1 or 1|0).

## Results

We compared Duet against four state-of-the-art SV callers, SVIM [12], cuteSV [11], NanoVar [13], and Sniffles [14], for SV calling and genotyping. We also compared Duet against LongPhase [24], an algorithm that can phase SVs based on an existing SV call set generated from other programs. We use the call set from cuteSV as the input for LongPhase. The benchmarking was on HG001, HG002, and HG00733, three standard human samples from the Human PanGenomics Project [25], with available high-confidence haplotype-resolved SV truth sets from HGSVC2 [18]. We tested the performance on three sequencing coverages: 8X (low-coverage), 20X (middle-coverage), and 40X (high-coverage). The summary results are presented in Fig. 2.

**Fig. 2.**
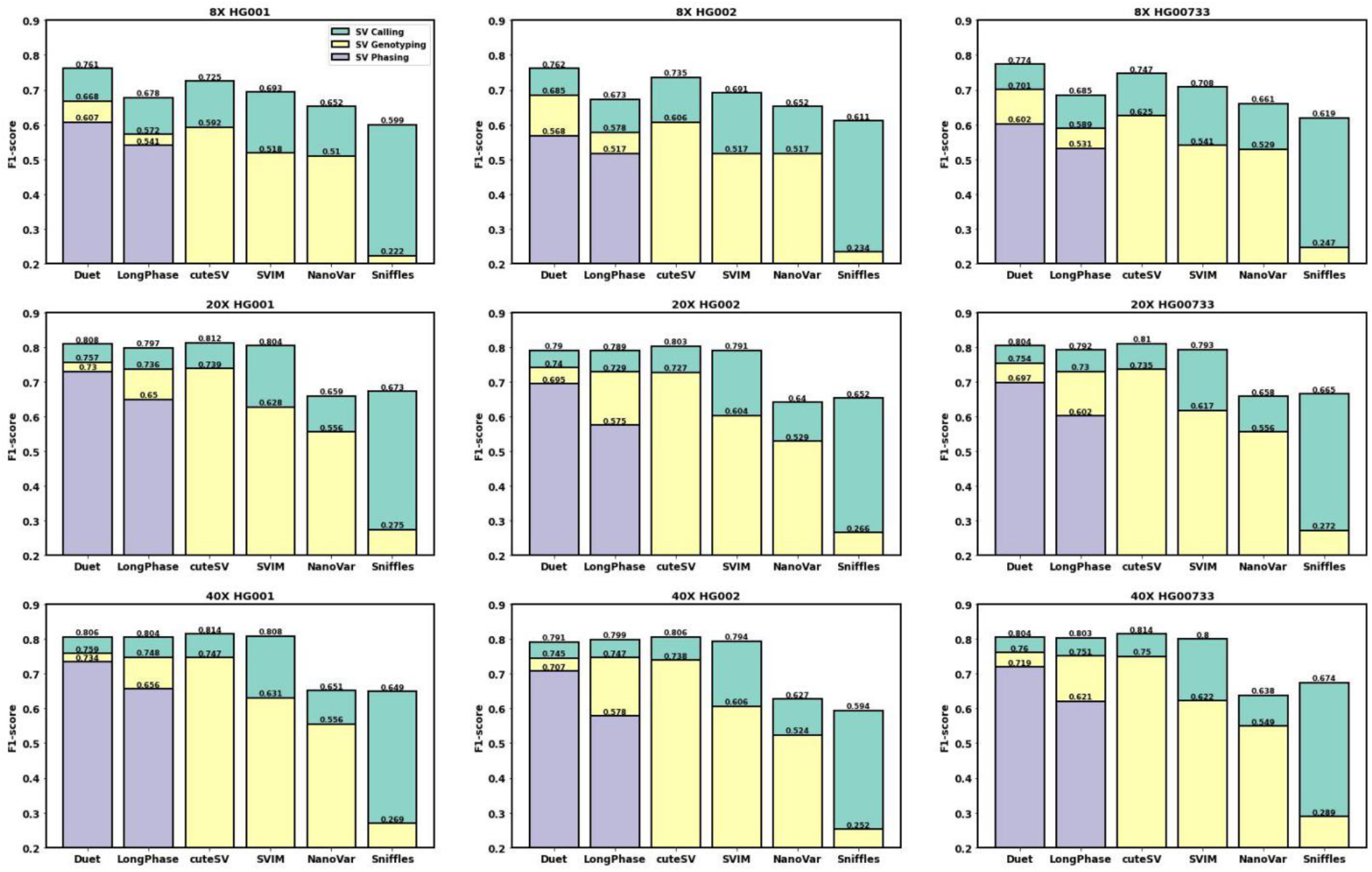
Benchmark results of SV detection tools for ONT data. The benchmark is on three different sequencing coverages across three human samples. Rectangles in green, yellow, and purple colors are performance of SV calling, genotyping, and phasing, respectively.

We have an integrated evaluation method for SV calling, genotyping, and phasing, with code available at [https://github.com/yekaizhou/duet/blob/main/src/scripts/evaluation.py]. The evaluation criteria of SV calling and genotyping is adapted from Truvari [https://github.com/ACEnglish/truvari].

In the evaluation of SV calling, an SV candidate is determined as a true positive (TP) if it meets the following conditions:

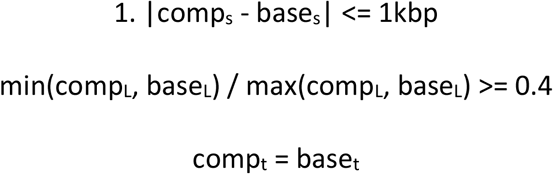

where comp_s_, comp_L_ and comp_t_ are the start coordinate, length, and the class of an SV prediction, respectively, while the base_s_, base_L_ and base_t_ are the starting coordinate, length, and the class of an SV recorded in the truth set, respectively. An SV call is counted as a false positive (FP) if it does not satisfy Eq. 1. A ground truth SV is assigned as a false negative (FN) if there is no SV call that satisfies Eq. 1 with it.

With the above definition, precision (or the ratio of TPs to total calls in predictions) is defined as:

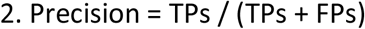

Recall (or the ratio of TPs to total calls in the truth set) is defined as:

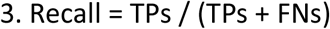

F1 score is defined as:

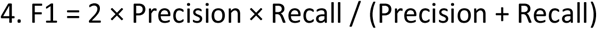

When considering the performance of SV genotyping, each TP SV call in SV calling gets further evaluated if its genotype is the same as the corresponding base call. If so, it is assigned to be a TP call, otherwise, it will be assigned as an FP call. All the FP calls in the SV calling evaluation procedure remain FP in SV genotyping. Each of the ground truth SVs will be evaluated and assigned TP if there is at least one TP call corresponding to it, otherwise it is assigned false negative (FN). Eqs. 2–4 are used to calculate the statistics of SV genotyping.

Switch error rate is a standard metric to evaluate the phasing accuracy of abundant and adjacent genome variants such as SNPs. However, this metric fails to be a reasonable metric for SVs, which are large and distant [24]. Therefore, we adapted and adjusted the standard evaluation method of SV genotyping for SV phasing. SV phasing will separately evaluate every phase block produced by WhatsHap: For each phase set, the haplotype prediction of TP SV calls in SV calling will have two versions, one remains original and the other was flipped (0|1 to 1|0; 1|0 to 0|1; 0|0 and 1|1 remains unchanged), and their haplotypes will be separately compared to the ground truth set: If the haplotype matches, it will be a TP call, otherwise, it will be an FP call. All the FP calls in the SV calling evaluation procedure remain FP in SV phasing. The evaluation will record the version with more TPs for every phase set. Each of the ground truth SVs will be evaluated and assigned TP if there is at least one TP call corresponding to it, otherwise it is assigned false negative (FN). Eqs. 2–4 were used to calculate the statistics of SV phasing.

At 8X sequencing coverage of the three human samples, Duet achieved up to 0.82 precision, 0.73 sensitivity, and 0.77 F1-score genome-wide for SV calling; 0.75 precision, 0.66 sensitivity, and 0.70 F1-score for SV genotyping; and 0.65 precision, 0.57 sensitivity, and 0.60 F1-score for SV phasing. Duet outperformed other tools in SV calling, genotyping, and phasing at the low coverage of 8X. Duet could phase SVs within a genome range of 2.37 Mbp on average (6.49 Mbp N50 and 0.81 Mbp median length), which included up to 355 SVs per block in our tested samples.

The performance of Duet has stable improvement when the sequencing coverage is increased to 20X, and achieved up to 0.85 precision, 0.77 sensitivity, and 0.81 F1-score genome-wide for SV calling; 0.79 precision, 0.72 sensitivity, and 0.76 F1-score for SV genotyping; and 0.76 precision, 0.70 sensitivity, and 0.73 F1-score for SV phasing. Duet showed increased improvement compared to other tools in SV phasing than the performance at 8X coverage. Duet still outperformed other tools in SV genotyping but LongPhase and cuteSV were quite close. Duet fell marginally short of cuteSV’s capability in SV calling.

The performance of Duet has slight improvement when the sequencing coverage is increased from 20X to 40X, and achieved up to 0.86 precision, 0.77 sensitivity, and 0.81 F1-score genome-wide for SV calling; 0.81 precision, 0.72 sensitivity, and 0.76 F1-score for SV genotyping; and 0.77 precision, 0.70 sensitivity, and 0.73 F1-score for SV phasing. Duet still has a great competing edge in SV phasing. As with 20X coverage, Duet outperformed other tools in SV genotyping for most samples although results from LongPhase and cuteSV are comparable. Similarly, Duet fell slightly short but comparable to cuteSV in SV calling.

## Discussion

In this study we developed a tool for SV calling, genotyping, and phasing, which we named Duet. We demonstrated that Duet is more accurate and powerful than other state-of-the-art tools in resolving genomic SVs, particuarly those that lead to disease, when only low-coverage data is available, which is very useful in clinical settings and resource-constrained lab settings where sufficient genomic data coverage may not be available. At higher coverages, Duet’s performance in SV phasing is further improved in comparison to other competing tools used for the purpose, while maintaining a comparable performance in SV genotyping and SV calling. The benchmark results of Duet further demonstrate that by incorporating SNP calling, SNP phasing, read haplotagging, the tailored decision tree is beneficial to SV detection, especially in accurate SV phasing and the quality control for low-coverage SV calling.

However, the benchmark results also indicate there are still some limitations in Duet:

I. Starting from middle-coverage, Duet becomes less competitive in SV calling. The main reason is that when the sequencing coverage goes higher, the feature p_SV_ used by other tools has recovered its discriminative power to remove false signals. Duet’s current quality control method has clear improvement at low-coverage, and will generally be caught up by other state-of-the-art methods when the coverage goes higher.

II. The performance improvement of Duet from middle-coverage to high-coverage is moderate. All the benchmarked tools have similar situation. This is because our highly accurate truth sets are derived from multi-platform sequencing, and some genome regions are hard to map using single-strand sequencing technology, even with high coverage. Despite this objective explanation, there is possibly room to improve the performance of Duet at high-coverage data by updating the quality control method to fit both low- and high-coverage sequencing data in a coverage-dependent manner.

## Conclusions

In this study, we introduced Duet, a long-read-based tool for accurate SV calling and phasing. Duet showed promising results that the incorporation of SNP signatures will largely boost the performance of SV detection, especially when there is constraint in sequencing coverage and when there is a need for accurate SV phasing. The tool’s great adaptability and scaling performance in SV calling and phasing promise its usefulness in routine clinical applications.

## Availability and requirements

**Project name**: Duet

**Project home page**: https://github.com/yekaizhou/duet

**Operating system(s)**: Platform independent

**Programming language**: Python

**Other requirements**: All software requirements are listed in https://github.com/yekaizhou/duet#dependencies

**License**: BSD-3-Clause

**Any restrictions to use by non-academics**: None

## List of abbreviations

FP: false positive
MAF: minor allele frequency
ONT: Oxford Nanopore Technologies
SNP: single-nucleotide polymorphism
SV: structural variant
TP: true positive
WGS: Whole genome sequencing

## Declarations

### Availability of data and materials

The source code of Duet and the scripts for data preparation and benchmarking are available at https://github.com/yekaizhou/duet. The hg38_NoALT human reference genome is available at: http://ftp.1000genomes.ebi.ac.uk/vol1/ftp/data_collections/HGSVC2/technical/reference/20200513_hg38_NoALT/hg38.no_alt.fa.gz. The HG001, HG002, and HG00733 sequencing reads datasets can be downloaded from: https://s3-us-west-2.amazonaws.com/human-pangenomics/NHGRI_UCSC_panel/HG001/nanopore/Guppy_4.2.2/, https://s3-us-west-2.amazonaws.com/human-pangenomics/NHGRI_UCSC_panel/HG002/nanopore/Guppy_4.2.2/, https://s3-us-west-2.amazonaws.com/human-pangenomics/NHGRI_UCSC_panel/HG00733/nanopore/Guppy_4.2.2/. The corresponding ground truth set for these three samples can be downloaded from: http://ftp.1000genomes.ebi.ac.uk/vol1/ftp/data_collections/HGSVC2/release/v2.0/integrated_callset/variants_freeze4_sv_insdel_sym.vcf.gz.

### Author’s contributions

Y.Z. developed the tool and wrote the manuscript. Y.Z. and A.W.S.L. benchmarked the Duet against the other tools. A.W.S.L., S.S.A., T.L., and R.L. revised the manuscript. R.L. conceived the project. All authors contributed to and approved the manuscript.

